# Counting on AR: EEG responses to incongruent information with real-world context

**DOI:** 10.1101/2024.08.22.608951

**Authors:** Michael Wimmer, Alex Pepicelli, Ben Volmer, Neven ElSayed, Andrew Cunningham, Bruce H. Thomas, Gernot R. Müller-Putz, Eduardo E. Veas

**Author notes:** Correspondance: Gernot R. Müller-Putz.

## Abstract

Augmented Reality (AR) technologies enhance the real world by integrating contextual digital information about physical entities. However, inconsistencies between physical reality and digital augmentations, which may arise from errors in the visualized information or the user’s mental context, can considerably impact user experience. This study characterizes the brain dynamics associated with processing incongruent information within an AR environment. We designed an interactive paradigm featuring the manipulation of a Rubik’s cube serving as a physical referent. Congruent and incongruent information regarding the cube’s current status was presented via symbolic (digits) and non-symbolic (graphs) stimuli, thus examining the impact of different means of data representation. The analysis of electroencephalographic (EEG) signals from 19 participants revealed the presence of centro-parietal N400 and P600 components following the processing of incongruent information, with significantly increased latencies for non-symbolic stimuli. Additionally, we explored the feasibility of exploiting incongruency effects for brain-computer interfaces. Hence, we implemented decoders using linear discriminant analysis, support vector machines, and EEGNet, achieving comparable performances with all methods. The successful decoding of incongruency-induced modulations can inform systems about the current mental state of users without making it explicit, aiming for more coherent and contextually appropriate AR interactions.

## 1. Introduction

Augmented Reality (AR) technologies enrich the physical environment by visualizing complementary digital information in real-time, thereby overcoming its natural limitations [1,2]. They can effectively communicate relevant data to users or provide semantic context to physical objects by overlaying situated information, such as symbolic and non-symbolic numbers [3]. Symbolic numbers, like numerical digits, offer precise and easily understood descriptions of the real world, whereas non-symbolic numbers, like graphs or quantities, foster the comprehension of proportional relations [4,5]. However, the presented information might not match users’ expectations, e.g., due to the users’ current mental context or errors in the underlying data yielding implausible information. Such perceived or factual incongruencies between the physical world and digital augmentations can impair the user experience. Possible strategies to address such issues include human-centric approaches [6], for example, detecting the processing of incongruent information utilizing physiological data from the user such as non-invasively recorded electroencephalography (EEG) signals.

The N400 was described as the “electrophysiological sign of the ‘reprocessing’ of semantically anomalous information” [7]. This component of event-related potentials (ERPs) derived from stimulus-related brain activity was first studied in language comprehension tasks and found after semantically incongruent sentence endings (“He spread the warm bread with *socks*.”). Even though the N400 was initially considered specific to language processing, it has been observed for numerous meaningful stimuli, including pictures [8], environmental sounds [9], and mathematical equations (“7 × 8 = *54*”) [10]. These components share various commonalities, e.g., in waveform and time course [11]. Additionally, there are intriguing functional similarities, i.e., the amplitude of the negativity is sensitive to semantic priming and context, even across modalities. For example, spoken words preceded by unrelated sounds elicit an N400 compared to those following related sounds [12]. Likewise, the N400 was found after the presentation of visual stimuli following incongruent olfactory primes [13,14]. Functional and temporal similarities of the N400 following incongruencies across different stimulus types and modalities support the idea of a common network responsible for semantic knowledge [15].

However, differences in topographical patterns of the EEG responses provide counterevidence. For instance, visually presented stimuli manifest in centro-parietal maxima [16], whereas the auditive N400 is more centrally distributed [17]. Other neuroimaging techniques, like magnetoencephalography, consistently identified temporal areas, e.g., the superior temporal gyrus [18,19] as a source of the N400 caused by semantic (linguistic) incongruencies. Interestingly, increased activity in the parietal angular gyrus was found after unrelated sounds and speech utilizing visual primes [20], suggesting a possible involvement in semantic integration from different meaningful stimulus types and modalities [21]. This accumulated evidence indicates that semantic knowledge is distributed across functionally similar but distinct brain areas whose involvements vary depending on the type of stimulus [15].

Despite the impact of N400 studies, e.g., on the understanding of language processing or semantic memory, there have been limited attempts to decode semantic incongruencies via brain signals. In clinical applications, the N400 has been used to analyze residual cognitive functions in non-communicating individuals or patients with disorders of consciousness [22]. For example, lexico-semantic processing was assessed in a picture-speech mismatch task and was detected in 50% of the participants via multivariate pattern analyses and support vector machines (SVMs) [23]. Disregarding the insufficient detection rate, EEG-based N400 decoding has the potential to offer an objective evaluation measure [24].

For more general applications, incongruency decoding can inform brain-computer interfaces (BCIs) [25] about the mental context of a user [26]. In prior work, participants were primed through visually presented words [27,28] or sentences [29] to modulate their responses to the following related and unrelated stimuli. Unrelated word pairs could be detected on a single-trial level with varying accuracies of 54 to 67% using logistic regression [28], and with an average accuracy of 58% in an extension of this experiment [27]. Similar performances were reported for the single-trial decoding of semantic violations in speech for different classification methods, i.e., SVM, random forest, and multilayer perceptron [29]. Notably, various ways of priming the mental context have been studied, including physical objects. For instance, the N400 could be detected following the presentation of words unrelated to an object held by the participants [26], comparable to the physical referent used in the current study. We refer to Dijkstra et al. [30] for an overview of BCI research exploiting the N400 effect.

In this work, we combine emerging AR technologies to present situated information with EEG-based decoding of incongruencies relevant to the design of BCIs. Particularly, we implemented an interactive and intuitive task featuring the manipulation of a Rubik’s cube as physical reference and context, as well as visualization of related information through symbolic and non-symbolic numbers. Hence, this paradigm represents a fundamental application area for AR by providing participants with complementary data about their physical surroundings. Successful detection of incongruent or implausible data could allow systems to react accordingly, e.g., by offering additional contextual information, explanations, or alternative options. Subsequently, we focus on the analysis of neurophysiological activity corresponding to the following research questions (RQs):

**(RQ1)** Which EEG responses correlate with processing incongruent information in AR related to a physical referent?

**(RQ2)** Are there differences in the responses to symbolic and non-symbolic information?

Based on these analyses, we additionally assess the feasibility of decoding incongruency effects for BCIs employed within AR interactions:

**(RQ3)** How reliably can the processing of incongruent information be detected from single-trial EEG?

## 2. Methods

### 2.1. Participants

Twenty participants (28.2 ± 6.2 years old (mean (M) ± standard deviation (SD)), 14 male, 6 female) took part in this study. All participants reported normal or corrected-to-normal vision, and none had a history of neurological or psychiatric disorders. Six participants self-reported having little or no experience with head-mounted displays (HMDs), and nine had no prior experience with EEG studies. The study received approval from the ethics committee of the University of South Australia and was conducted following the Declaration of Helsinki (1975). The participants signed informed consent and received vouchers worth $40 AUD.

### 2.2. Experimental setup and procedure

Before the recordings, the participants’ color vision was assessed through an Ishihara test (https://www.colorblindnesstest.org/ishihara-test/) [31]. After successful completion, they were comfortably seated at a table such that they could easily reach and grab a tricolor (red, blue, and white) Rubik’s cube (Fig. 1). A camera (Canon EOS 200D II, Tokyo, Japan) pointed at the cube to detect the nine colors that comprised its top surface. The color detection was implemented based on the Qbr Rubik’s cube solver (https://github.com/kkoomen/qbr). The AR visualizations were displayed using a HoloLens 2 HMD (Microsoft, Redmond, WA, USA) and designed in Unity 2021.1.31 (https://unity.com).

**Fig. 1:**
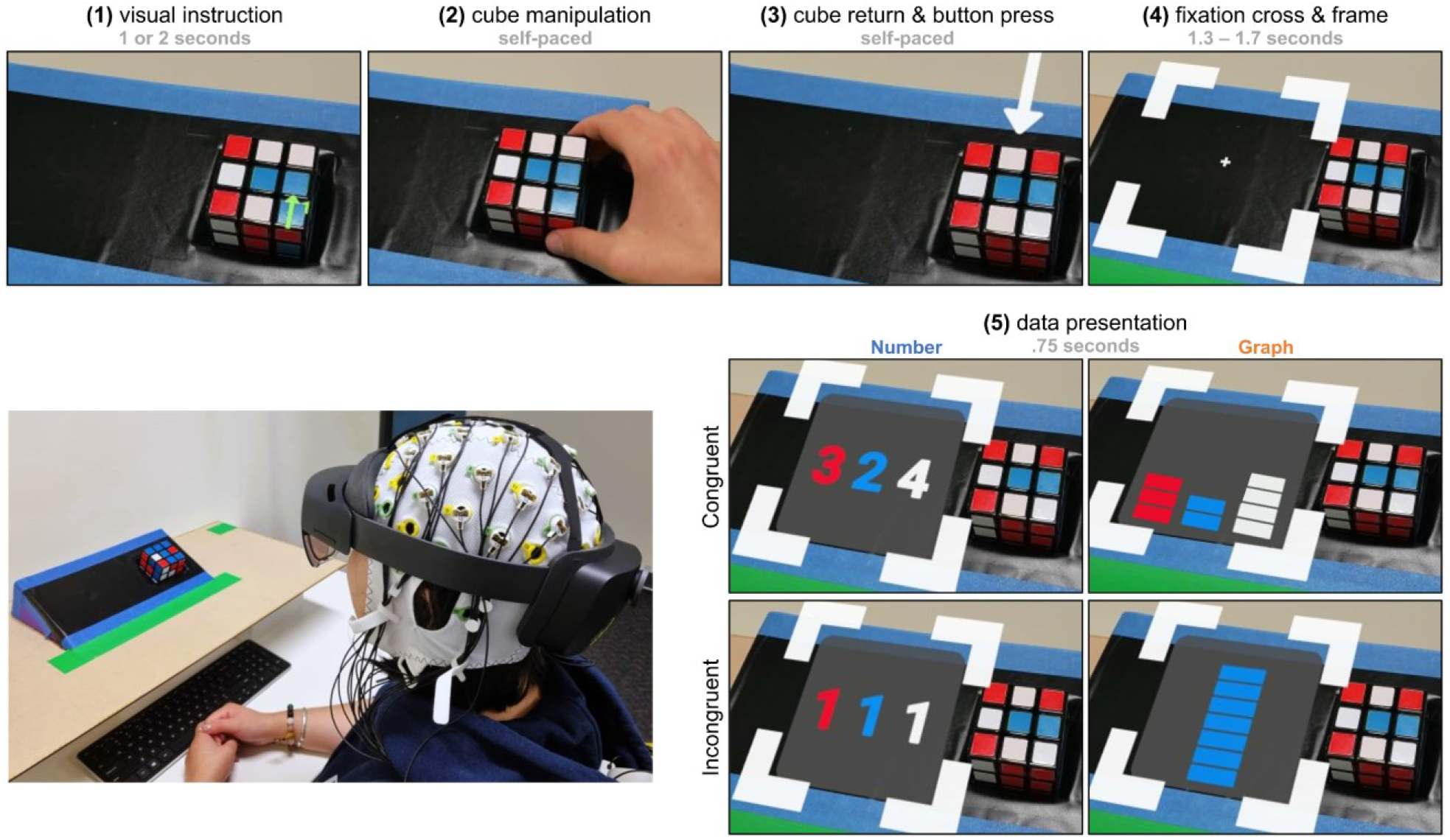
Experimental setup and timing of one trial. A participant wearing the EEG cap and the HoloLens 2. The Rubik’s cube is in its initial position. **(1)** Participants received manipulation instructions through visual cues. **(2)** After performing the indicated manipulations, participants placed the cube in its initial position. **(3)** Participants counted the number of red, blue, and white squares and pressed a keyboard button upon completion. In the pictured example, the count is 3-2-4. **(4)** Subsequently, a fixation cross inside a frame appeared for the following data presentation. **(5)** A congruent or incongruent count was presented inside the frame. Depending on the current run, the count was visualized using *Number*s or *Graphs*.

The implemented 2×2 experimental design consisted of two groups, i.e., symbolic (*Number*) and non-symbolic (*Graph*) data presentations, and two conditions, i.e., congruent and incongruent. Symbolic data presentations visualized information through Arabic numerals, while bar charts were used for non-symbolic presentations. The experimental protocol encompassed 12 runs of 33 trials (23 congruent, 10 incongruent) following a randomized order. Runs were alternated between those featuring *Number*s and *Graph*s, e.g., *Number, Graph, Number*, et cetera. Half of the participants commenced the experiment with *Numbers* and the other half with *Graphs*. Participants took self-paced breaks of usually one to five minutes between runs to prevent fatigue. Beforehand, participants performed one training run of 10 trials of each group to familiarize themselves with the paradigm. After the final run, participants completed a questionnaire.

Figure 1 illustrates the experimental setup and the timing of one trial. At the beginning of each trial, participants were instructed on manipulating the cube using one or two visual cues, i.e., green arrows. They were presented for 1 or 2 seconds (s) and indicated the rows and columns for rotation in the given directions. Upon manipulation, participants restored the cube to its initial position. Next, the participants counted the number of red, blue, and white squares on the cube’s top surface and indicated they knew the correct count by pressing a physical keyboard button. This initiated the presentation of a fixation cross inside a frame left to the cube. At this point, participants were asked to concentrate their gaze on the cross for the rest of the trial to minimize ocular artifacts. After 1.3 to 1.7 s (randomized), a congruent or incongruent count, informed by the color detection, was displayed inside the frame for 0.75 s following the order red-blue-white. Congruent counts matched the colors currently constituting the cube’s top surface, while incongruent counts were inherently impossible, e.g., 0-7-0 or 2-1-1. We prepared a set of 21 different incongruent counts to prevent accustoming to stimuli. A pause of 0.75 s followed each data presentation before a new trial commenced with a countdown from two to zero (1.5 s).

A video of the experiment can be found in the supplemental material. The experimental paradigm is available at https://github.com/WearableComputerLab/RubiksAR.

### 2.3. Data acquisition

To record EEG signals, we used an ActiCap system connected to a BrainAmp amplifier (Brain Products, Gilching, Germany). We positioned 32 electrodes according to the international 10-10 system at AFz, F3, F1, Fz, F2, F4, FC5, FC3, FC1, FCz, FC2, FC4, FC6, C5, C3, C1, Cz, C2, C4, C6, CP5, CP3, CP1, CPz, CP2, CP4, CP6, P3, P1, Pz, P2, and P4. The ground and reference electrodes were placed at Fpz and the right mastoid, respectively. The data was sampled at 500 Hz. Before the recordings, we ensured the impedances between the scalp and the electrodes were below 10 kΩ. We further monitored the EEG signals throughout the experiment.

The HMD was used to record gaze data at a sampling frequency of 30 Hz during the stimulus presentation. EEG and gaze signals were synchronized with the event markers of the experimental paradigm utilizing the Lab Streaming Layer framework [32].

### 2.4. Data preprocessing

The data processing and analysis were performed in Matlab R2022a (The MathWorks, Natick, MA, USA) utilizing functions provided in the EEGLAB toolbox 2022.0 [33] and the BioSig toolbox [34], and Python 3.9.12.

EEG signals were bandpass filtered between 1 and 25 Hz (Butterworth, 4th order, zero-lag). To suppress noise stemming from the HMD and the power line, we applied notch filters at 30 Hz and 50 Hz (both Butterworth, 2nd order, zero-lag). Next, we resampled the data to 125 Hz to reduce the computational effort and performed independent component analysis applying the extended infomax algorithm [35]. We visually inspected the components to remove those contaminated with artifacts from eye movements or blinks. Subsequently, we segmented the continuous signals into epochs of 5 s, i.e., -2 s before to 3 s after the data presentation. At this point, an average of 194 trials were available per participant and group for further analysis.

We rejected epochs based on visual inspection and objective measures, i.e., amplitude threshold (exceeding ± 35 µV), kurtosis, and joint probability (both 5 ⋅*SD*) [36]. Further, we rejected epochs with excessive eye movements. For each participant and epoch, we computed the variance of the Euclidean distance between the gaze positions and the fixation cross and rejected epochs with a z-score outside ± 3. On average, we rejected 9% of the trials in both groups. Finally, we spherically interpolated 1.3 ± 1.2 (*M* ± *SD*) noisy channels per participant based on visual inspection. One participant could not identify incongruent counts and was therefore excluded from the analysis.

### 2.5. Time-frequency analysis

We identified event-related desynchronizations and synchronizations (ERD/S) [37,38] to study the time-frequency behavior in both groups and conditions. ERD/S was calculated by bandpass filtering all event-related epochs, computing the power of individual samples by squaring their amplitudes, and averaging over epochs [39]. Before squaring, we subtracted the mean signal to suppress evoked potentials that could mask the induced activity [40]. ERD/S is defined as a power decrease (ERD) or increase (ERS) during the data presentation *A* relative to a reference period *R*, i.e., [-0.5,0] s before the stimulus onset. We computed ERD/S values for frequencies between 4 and 25 Hz with a bandwidth of 2 Hz according to *ERD/S* (%) *= (A-R)/R* ⋅100 [41]. The statistical significance of ERD/S values was assessed using a non-parametric *t*-percentile bootstrap algorithm [39,42] with a significance level of α = 0.05. These time-frequency decompositions (or ERD/S maps) were calculated for each participant.

Subsequently, we computed grand average ERD/S maps for each group and condition by averaging the 19 individual maps. To include the statistical significance at the participant level, we set all non-significant ERD/S values to zero. Further, we only considered event-related changes of at least 5% in the grand average maps [43]. Additionally, we derived ERD/S for the theta (4 - 8 Hz), alpha (8 - 12 Hz), and beta (12 - 25 Hz) frequency bands separately. Finally, we computed the maximum ERD/S values for theta and the minimum values for alpha and beta within [0.35,0.7] s after stimulus onset for further analyses.

### 2.6. Binary classification approach

We decoded incongruent trials in both groups using three classification approaches, i.e., (i) shrinkage linear discriminant analysis (sLDA) [44], (ii) SVMs, which are both commonly used in ERP classification problems [45], and (iii) EEGNet, a compact convolutional network that has been successfully employed in a variety of BCI paradigms [46]. The binary classifications were performed using the preprocessed data, as described in Section 2.4.

#### 2.6.1. Classification with sLDA and SVM

We decoded congruent and incongruent trials in both groups separately. For each participant, we derived time domain features from amplitude values and statistical measures. Specifically, for each channel, we computed the mean of overlapping windows of 152 ms in steps of 32 ms [47], approximately based on the grand average ERPs. The first window started at *t* = 0.3 s relative to the stimuli onset, and the last window at *t* = 0.85 s, e.g., [0.3,0.452] s, [0.332,0.484] s, et cetera. This resulted in 18 features per channel. The standardized features were used to train sLDA classifiers and SVMs with a linear kernel as the basis function. In particular, we trained and tested the personalized models via a stratified 10 times 5-fold cross-validation (CV) approach. These binary classification problems consisted of balanced classes by randomly choosing a different subset of congruent trials equal to the number of incongruent trials in each of the 10 iterations. The reported accuracies are averages of 50 folds per participant (5 folds times 10 iterations).

#### 2.6.2. Classification with EEGNet

To assess the utility of deep learning approaches for decoding incongruency, we used the EEGNet architecture proposed by Lawhern et al. [46]. EEGNet was designed to classify EEG data in sequential convolution steps. First, temporal convolutions are performed with *F*_*1*_ 2D convolutional filters that mimic bandpass frequency filters with kernel weights identified from the EEG data. Next, depthwise convolutions learn *D* spatial filters for each temporal filter to integrate information regarding channel locations. The depthwise convolutions reduce the number of trainable parameters to fit as they are not fully connected to the previous outputs. Further, a separable layer consisting of depthwise convolutions followed by *F*_*2*_ = *F*_*1*_ ⋅*D* pointwise convolutions combines information across filters more efficiently. Each convolution layer is followed by batch normalization, average pooling, and dropout layers to mitigate overfitting. Finally, flatten, dense, and softmax layers generate the classification output. We refer to the original work for an exhaustive description of the architecture.

For our application, we learned 16 spatiotemporal filters (*F*_*1*_ = 8, *D* = 2) with a temporal kernel size of 32. The size of the depthwise convolution equals the number of EEG channels (32). The models were trained for 500 iterations with a minibatch size of 64 using categorical cross-entropy as the loss function and Adam as the optimizer. The dropout rate was set to 0.5 for all layers [48]. As input features, we used all amplitude values of the EEG signals within [0.3,0.85] s relative to the stimulus onset. The CV approach with balanced classes is analogous to Section 2.6.1 and closely resembles the methodology presented in the original work [46].

### 2.7. Statistics

We performed sample-wise Wilcoxon signed-rank tests (α = 0.05) to compare ERPs at CPz (Fig. 3). Similarly, we performed Wilcoxon signed-rank tests (α = 0.05 and 0.01, respectively) to find electrodes with significantly different responses to incongruent and congruent stimuli at the time points given in Fig. 2. The same tests were further used to assess differences (α = 0.05) in (i) latency and amplitude of the N1 component of ERPs elicited by *Numbers* and *Graphs* (Fig. 4), (ii) minimum and maximum ERD/S values within [0.35,0.7] s for theta, alpha, and beta frequency bands (α = 0.05), and (iii) subjective ratings obtained in questionnaires after the experiment (Fig. 7). Finally, we performed a Friedman test (α = 0.05) for each group to compare the decoding performance of the implemented classification methods (Fig. 6). We applied the false discovery rate (FDR) procedure [49,50] to correct for multiple testing. Non-parametric methods were preferred due to the small sample size (*N* = 19) and to mitigate influences by outliers [51–53].

**Fig. 2:**
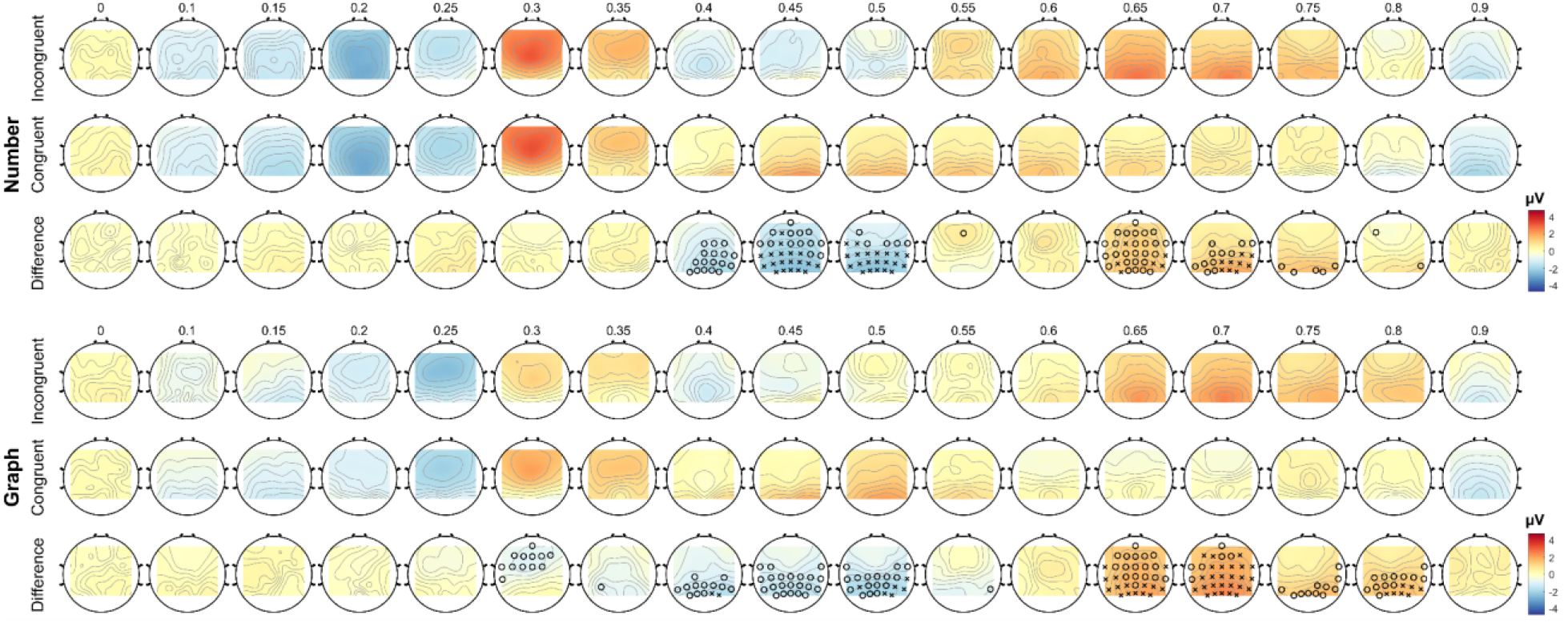
Topographical distribution of the grand average ERPs. Shown are the topographies for Number (top) and Graph (bottom) at the time points indicated above the respective top rows (in s, relative to the stimulus onset at 0 s). Difference plots (bottom rows) are obtained by subtracting congruent (middle rows) from incongruent (top rows) signals. Electrodes with significant differences (Wilcoxon signed-rank, FDR corrected) are indicated with *o* (*p* < 0.05) or *x* (*p* < 0.01).

**Fig. 3:**
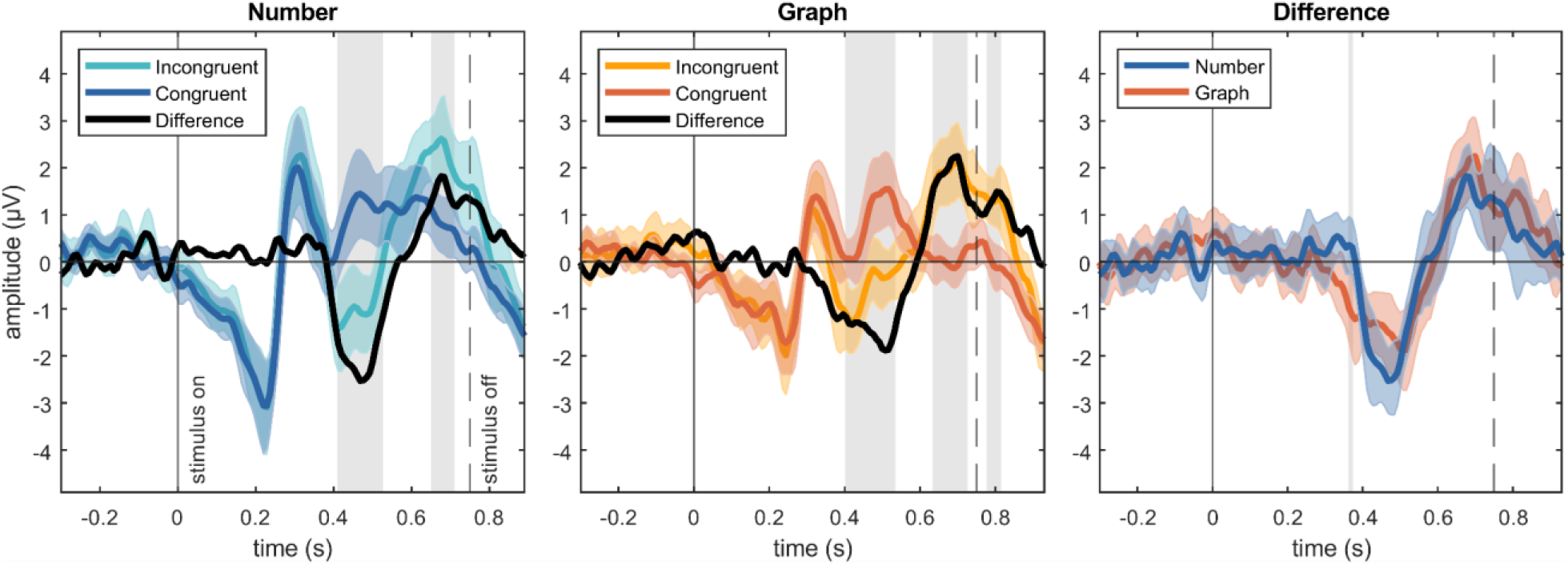
Grand average ERPs at CPz. Shown are the ERPs resulting from the processing of congruent (darker colors) and incongruent (lighter colors) stimuli for *Number* (left) and *Graph* (middle). The dashed lines indicate the differences *incongruent* - *congruent*, which are also shown in the right subfigure. The shaded areas indicate ± 2 ⋅SEM. The gray segments show significant differences (Wilcoxon signed-rank, FDR corrected, *p* < 0.05). All timings are relative to the stimulus onset at 0 s.

**Fig. 4:**
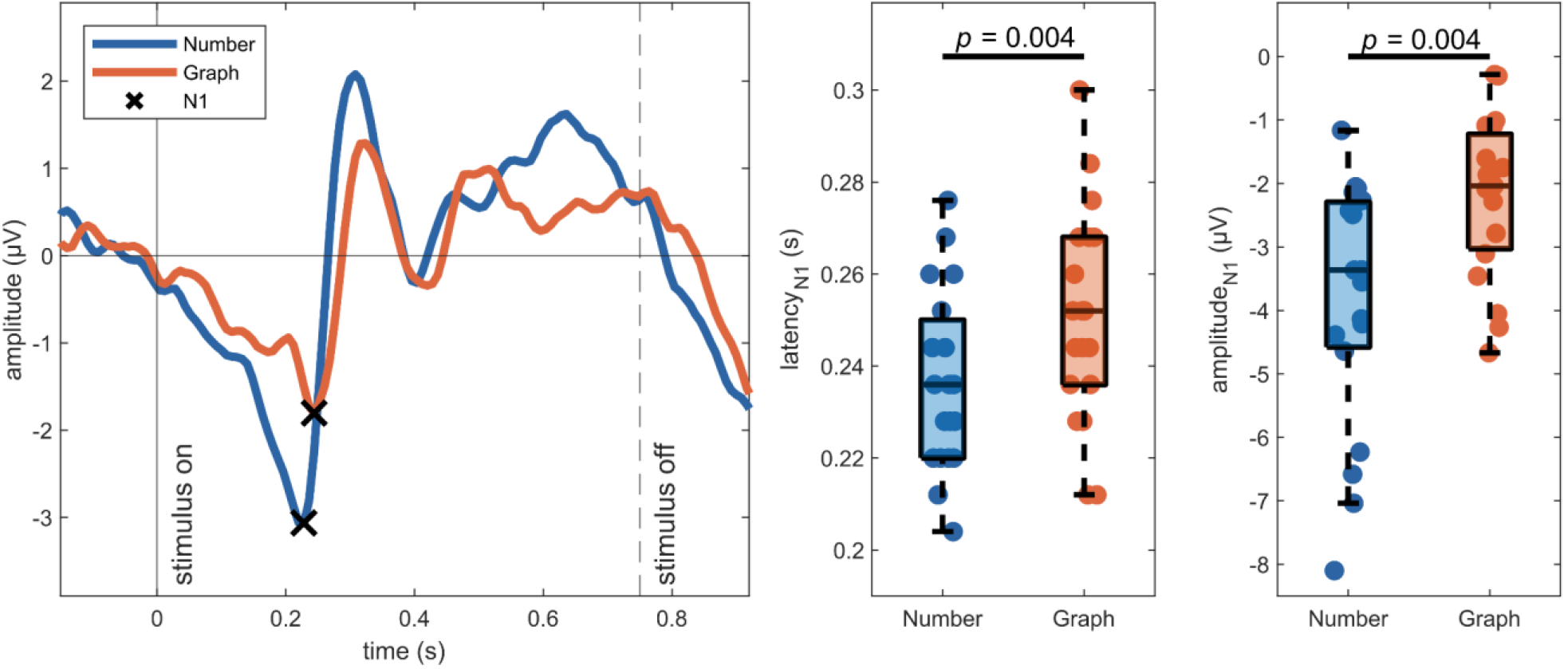
Comparison of latency and amplitude in responses to symbolic and non-symbolic stimuli. Shown are the ERPs for *Number* and *Graph* (congruent and incongruent combined). The first negative peaks N1 are marked. The participant-level distributions of the latency and amplitude of N1 are illustrated. The results of the statistical tests are given (Wilcoxon signed-rank, FDR corrected). All timings are relative to the stimuli onset at 0 s.

The individual significance levels [54] used to assess the classification results (Fig. 6a) were obtained using cumulative binomial distributions as expressed in Eq. (1) [55]. The probability *P* of predicting a condition correctly *k* times is defined as

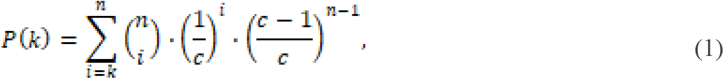

where *n* is the number of trails, and *c* = 2 is the number of conditions. We present the individual significance thresholds per participant and group obtained at α = 0.05.

## 3. Results

### 3.1. Incongruent information elicits N400 and P600 components

In Fig. 2, we show the topographical distributions of the ERPs following the presentation of congruent and incongruent information (**RQ1**). In both conditions, i.e., *Number* and *Graph*, we found incongruencies to cause more negative responses starting at approximately 400 ms after stimulus onset peaking in centro-parietal areas of the cortex. At around 650 ms, topographies show a centro-parietal positivity elicited by incongruent stimuli. Electrodes exhibiting significant differences in the given time points, i.e., 0 to 0.9 s after stimulus onset, are marked with *o* (*p* < 0.05) or *x* (*p* < 0.01). The differences *incongruent - congruent* have a slight bias toward the right hemisphere.

Given incongruencies manifest in centro-parietal cortical areas, we investigate the ERPs at CPz in greater detail in Fig. 3. For both conditions, we show the EEG correlates of incongruent (darker colors) and congruent (lighter colors) stimuli (± 2 ⋅standard error of the mean (SEM)). The differences *incongruent - congruent* are shown in black (dashed) and in the right subfigure. Statistically significant time points are indicated in gray.

For *Number* (left), the difference signal shows an initial negative peak at 0.468 s (−2.53 µV), followed by a positive peak at 0.676 s (1.82 µV). These components differ significantly in the segments [0.412,0.532] s and [0.652,0.708] s after stimuli presentation. Incongruencies in *Graph* (middle) cause a negative component peaking at 0.516 s (−1.87 µV) and a positive peak at 0.700 s (2.24 µV). Significant differences were found in the segments [0.404,0.532] s, [0.636,0.724] s, and [0.780,0.812] s. Incongruencies in symbolic and non-symbolic stimuli cause similar difference signals (right) with statistical significance only at 0.364 s.

### 3.2. Non-symbolic numbers elicit delayed EEG responses

The ERPs for *Number* and *Graph* were obtained by averaging all epochs of the respective groups disregarding the condition (Fig. 4, left) (**RQ2**). The components of these ERPs differ in latency (middle) and amplitude (left). To exemplify this, we investigated both metrics for the initial negative peaks (N1) in more detail. The latency for the N1 for *Graph* at 0.244 s is significantly delayed (*p* = 0.004) to the N1 for *Number* at 0.228 s, i.e., by 16 ms. Similarly, statistical tests revealed a difference in amplitude between groups (*p* = 0.004). Latencies in further components show similar behavior. Likewise, the cross-correlation between the respective grand average ERPs peaks at 16 ms too. These results must be carefully analyzed by considering the influences of the experimental paradigm (see Section 4.1).

### 3.3. Time-frequency analysis reveals incongruency-related theta synchronizations

Figure 5 shows incongruency-induced desynchronizations and synchronizations in the theta, alpha, and beta frequency bands. Alpha and beta bands exhibit power suppressions after stimulus presentation. However, these do not feature significant differences for congruent and incongruent information. Additionally, we observed a band power increase in theta peaking at approximately 360 to 460 ms after stimulus presentation. The maximum theta ERS is significantly stronger for incongruent stimuli in *Number* (*p* = 0.038) and *Graph* (*p* = 0.023).

**Fig. 5:**
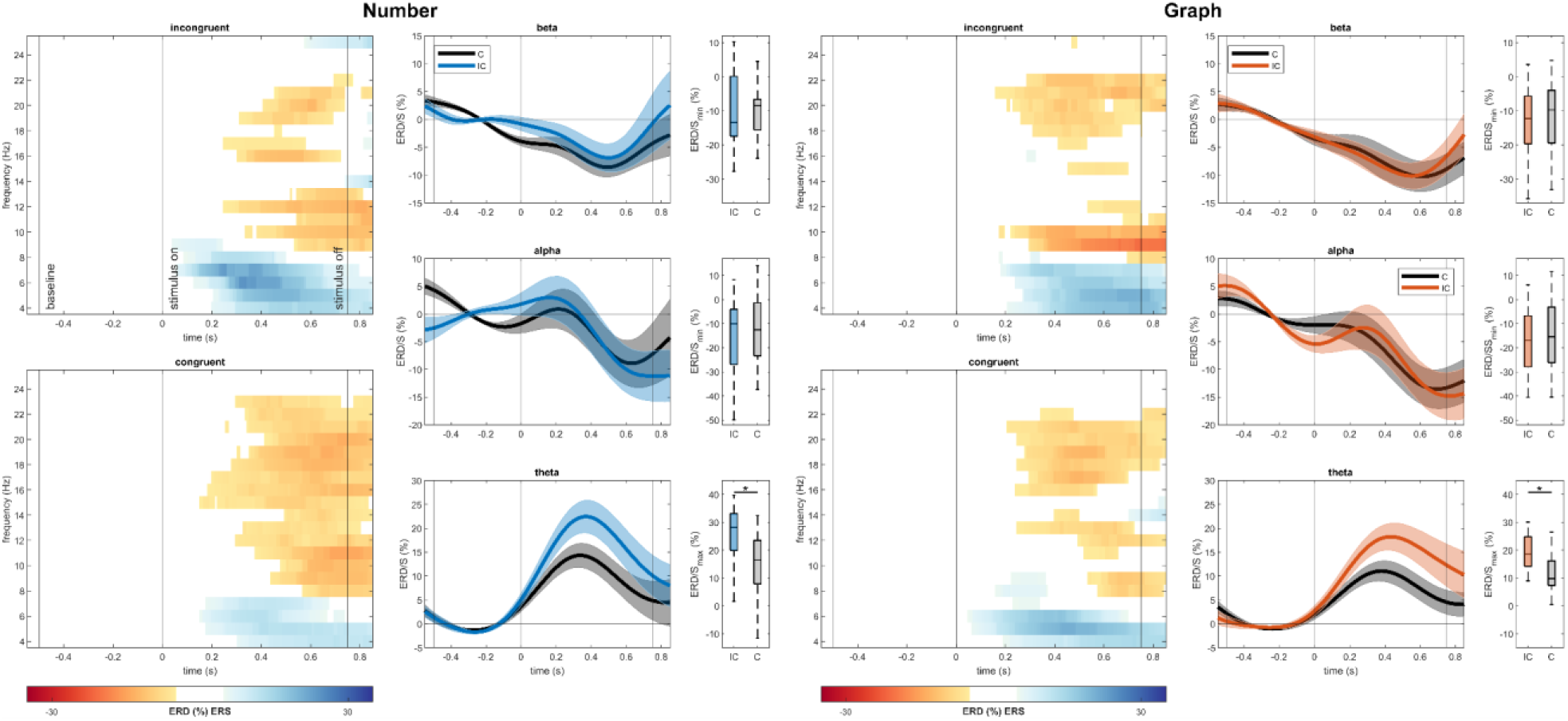
Grand average ERD/S results at CPz. Grand average ERD/S maps (left panels) for *Number* (left) and *Graph* (right) are shown for incongruent (IC) and congruent (C) stimuli. Red and blue colors indicate significant ERD and ERS, respectively (bootstrap, α = 0.05). Vertical lines mark the reference period ([-0.5,0] s), stimulus onset (0 s), and offset (0.75 s). The middle panels display the ERD/S for theta, alpha, and beta frequency bands (*M* ± *SEM*). The right panels show the corresponding distributions of the participant-level maximum (theta) and minimum (alpha, beta) ERD/S values within [0.35,0.7] s. Significant differences (Wilcoxon signed-rank, FDR corrected) are indicated (\**p* < 0.05).

### 3.4. sLDA, SVM, and EEGNet offer similar performance

Figure 6 depicts the distributions of the participant-level classification results for each group and classification method (**RQ3**). The Friedman test revealed no significant difference for *Number* (χ^2^(2) = 0.42, *p* = 0.810) or *Graph* (χ^2^(2) = 5.16, *p* = 0.114). For *Number*, 14 out of 19 participants (74%) achieved accuracies above their individual significance level with sLDA (classification accuracy: *M* = 63.8%, *SD* = 9.1%), while 16 (84%) did so with SVM (*M* = 64.4%, *SD* = 8.2%) and 15 (79%) with EEGNet (*M* = 64.8%, *SD* = 9.8%). Further, using *Graphs*, 16 participants achieved significant results with sLDA (*M* = 64.4%, *SD* = 5.8%) and SVM (*M* = 65.7%, *SD* = 6.7%), and 18 (95%) with EEGNet (*M* = 66.0%, *SD* = 6.9).

**Fig. 6:**
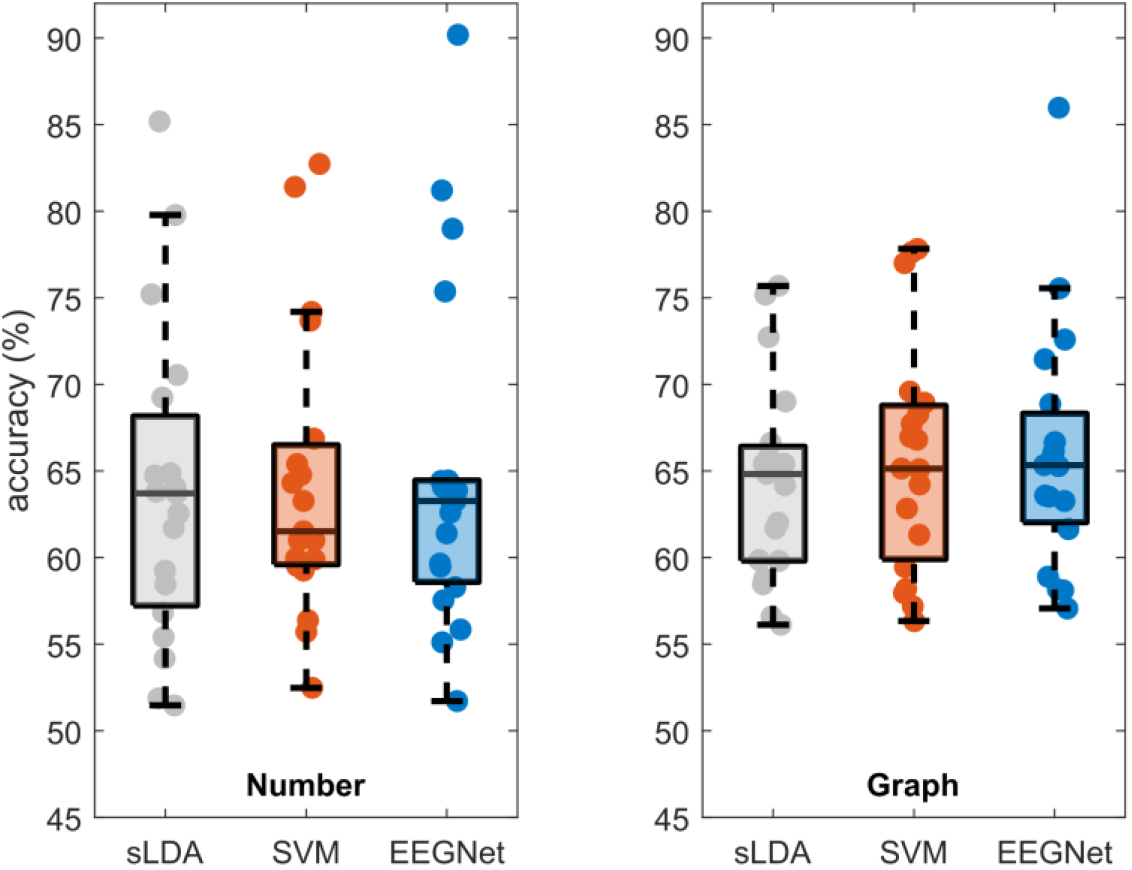
Classification results. Participant-level classification results for *Number* (left) and *Graph* (right) using sLDA (gray), SVM (orange), and EEGNet (blue). Statistical tests revealed no significant difference within either group (Friedman, FDR corrected, *p* < 0.05).

**Fig. 7:**
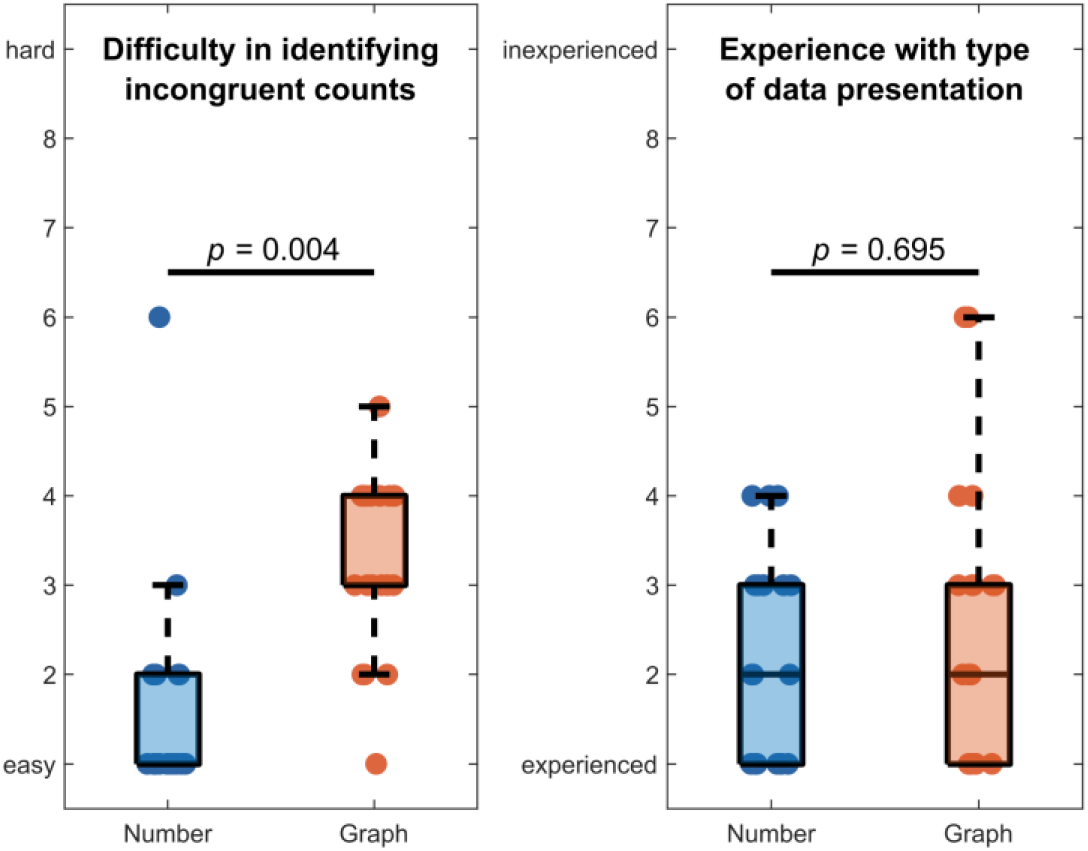
Subjective ratings. Answers from the post-experimental questionnaires. Participants were asked to assess (i) how difficult it was to identify incongruent counts using *Numbers* and *Graphs* (left) and (ii) how experienced they were using the respective means of data presentation (right). The results of the statistical tests are given (Wilcoxon signed-rank, FDR corrected).

### 3.5. Post-experimental questionnaires

Participants completed questionnaires concerning their subjective evaluations of using symbolic and non-symbolic numbers for presenting data. The analysis of these questionnaires revealed that participants found it significantly easier (*p* = 0.004) to identify incongruent counts presented with *Numbers* (*M* = 1.58, *SD* = 1.22) than with *Graphs* (*M* = 3.26, *SD* = 0.99). However, participants were comparably experienced (*p* = 0.695) in using *Numbers* (*M* = 2.26, *SD* = 1.15) and *Graphs* (*M* = 2.58, *SD* = 1.57). Participants could rate on scales from 1 (easy / experienced) to 9 (hard / inexperienced).

## 4. Discussion

The current work is among the first to study the effects of incongruency in AR environments. Hence, we carefully designed the experimental paradigm to obtain stimulus-related EEG responses unaffected by interferences, such as movement-related or ocular artifacts. Consequently, we divided the task into two distinct phases. Initially, participants actively engaged with the Rubik’s cube, after which they assumed a passive role during the stimulus presentation. This paradigm resembles a fundamental objective of AR technologies, i.e., providing users with in situ supplementary information about their physical environment [56,57]. As in this experiment, this information is typically derived from available data and does not necessarily match the users’ expectations [58], given their current mental context or the intrinsic implausibility of the presented information. Adaptive systems [59] could provide additional insights about the presented information, explanations regarding visualizations, or alternative options. Being able to recognize when an incongruency has been observed can further help inform the design of AR cues and raise the possibility of adaptive AR cues. One specific scenario is predictive AR cues for complex situations involving large groups of people interacting such as sports games, where prior work has investigated effective AR cues to represent predicted player trajectories based on deep learning models [60].

### 4.1. Neurophysiological results

The neurophysiological responses to incongruent information in this novel paradigm align with the existing literature on the N400 effect in monitor-based tasks (**RQ1**). Particularly, the observed ERPs exhibit a centro-parietal maximum, as commonly observed in related experiments using visual stimuli [61], including studies featuring numbers or mathematical problems [10,62,63]. As in these works, the spatial distribution of the ERPs presented in this paper is more pronounced in the right hemisphere. Given the bilateral but left-dominant localization of the neural generators, the right bias seems counterintuitive. Previously, it has been speculated that a “paradoxical lateralization” [12] attributed to tissue morphology accounts for the N400 topography.

Additionally, we observed a late positive component peaking just after 600 ms post-stimulus in centro-parietal brain areas. This finding was not entirely expected, as the P600 is commonly considered to be related to syntactic incongruencies in linguistic paradigms [64]. However, P600 components were reported in related studies, e.g., during arithmetic tasks [10,62,63,65], which contradicts this theory. It has been suggested that the P600 may reflect plausibility judgments or double-checking answers [62,63,66], and was even found to be sensitive to the degree of incongruency [65]. Likewise, similar patterns were reported in semantic violations in implausible verb-object relations [67]. For example, “Tyler canceled the *birthday…*” evoked only an N400 due to its implausibility, as opposed to “*subscription*”, whereas “Tyler canceled the *tongue…*” elicited both N400 and P600 components. The authors argued that these responses reflect the perceived impossibility of incongruencies. According to this line of reasoning, the P600 observed in the current study is linked to the plausibility evaluation for incongruent or impossible counts, such as 0-0-8 or 3-1-1. Notably, there are still unresolved discussions on the possibility that the P600 is a variant of the P3b subcomponent of the P300 [68]. Indeed, there are compelling similarities in time course, scalp distribution, and polarity. Moreover, both components have been observed in response to unexpected events, and their amplitudes are modulated by the stimulus probability [69]. Further research emphasizes the impact of task complexity and relevance, i.e., the P3b amplitude decreases with stimulus evaluation difficulty, while the latency increases [70,71], consistent with the reasoning of other P600 studies [10,72]. Hence, the late positivity in this work might reflect the responses to unlikely stimuli given that only 30% of the presented counts were incongruent. However, patient studies provided counterevidence. For instance, in one work with participants with basal ganglia impairment, ERPs did not reveal a P600 effect, while a P3b was elicited in an oddball paradigm [73]. Consequently, the authors argued that different neural sources are involved in generating these components, which would support their separability.

In addition to incongruency effects, we analyzed differences in the ERPs related to processing symbolic and non-symbolic stimuli (**RQ2**). The grand average responses show differences in latency and amplitude, i.e., *Graphs* elicited delayed ERPs with smaller amplitudes than *Numbers*. However, the experimental design influenced these results, as participants were instructed to focus their gaze on the fixation cross. The subsequent presentation of *Numbers* consistently appeared at this exact location, i.e., in their foveal field, whereas *Graphs* often encompassed only their peripheral field. For example, *Graphs* of common counts like 4-3-2 or 3-3-3 were visualized below their current gaze position (see Fig. 1 or supplemental video). Previous research found that visual stimulation of the foveal field evokes potentials with larger amplitudes than analogous peripheral field stimulation [74]. Consequently, the difference in amplitude does not necessarily reflect the cognitive processes involved in the stimulus evaluation but rather the stimulus site. The variations in latency are more informative. Neurophysiological evidence suggests that the speed of visual processing increases with eccentricity [75], hence the response to (peripherally presented) *Graphs* should be faster than *Numbers*. However, we observed the opposite, arguably indicating the additional efforts required to process the more complex non-symbolic stimuli [72]. In fact, in post-experimental questionnaires participants reported that identifying incongruent stimuli was significantly harder using *Graphs*. Additionally, there exists literature that found that humans are faster in comparing symbolic than non-symbolic numbers [76,77], which aligns with the results of this work.

Previous studies investigating band power modulations related to incongruency effects have primarily focused on language comprehension. The first evidence for theta band power increases was found following grammatical violations in sentence processing [78]. Subsequent research confirmed their results [79,80] and argued that an increased theta power reflects general working memory processes related to integrating information into a given context [81]. Indeed, theta synchronizations have been linked to elevated demands in working memory earlier [82,83], suggesting a broader functional role of low-frequency power modulations [84]. Further, they are not specific to linguistic paradigms but have also been observed, e.g., after visual stimuli [85,86]. Consequently, the theta ERS found in this paper is likely associated with the increased working memory load required for evaluating incongruent information.

### 4.2. Incongruency decoding for BCIs

Only a few works attempted single-trial decoding of incongruent stimuli using EEG signals (**RQ3**). Commonly reported decoding accuracies are just below 60% [27–29], compared to 66% achieved in this study. In a recent work on parts of this data set, we decoded incongruent *Numbers* with an average accuracy of around 63% [87]. Interestingly, the fact that participants found it significantly harder to identify implausible *Graphs* is not reflected in the classification results. Further, the classification results exhibit high variabilities between participants, similar to the findings of previous works that reported difficulties in extracting the N400 in individual participants [30]. Significant N400 effects are commonly reported to be found in the ERPs of approximately 50 to 70% of neurophysiologically healthy participants, with considerably lower success rates in individuals with disorders of consciousness. Having a proportion of potential end users that do not generate detectable brain patterns is not unique to N400 decoders but an issue in many BCI approaches [88]. Nevertheless, it limits the practical usability of such BCIs.

However, the classification accuracies of incongruency decoders have to be improved for real-world applications. The low signal-to-noise ratio of the N400 effect is a limiting factor. Consequently, a prior work investigated how the number of trials available for training impacts incongruency decoding [30]. With 400 trials per condition, the N400 could be classified above chance in 15 out of 19 participants, compared to only 6 out of 19 participants when recording 100 trials per condition. Deep learning approaches particularly profit from large amounts of data to facilitate high performance [89]. Another strategy to improve classification could be integrating multimodal physiological data into a hybrid BCI [90,91]. An increasing number of consumer-grade HMDs feature sensors and cameras to acquire additional signals likely improving BCI accuracies, e.g., pupil size or gaze [92]. In the current study, we recorded eye movement data for artifact rejection. However, we could not utilize them for classification after instructing participants to fixate their gaze on the cross during the stimulus presentation. Hence, while minimizing the influence of ocular artifacts on the EEG responses, we could not analyze potential differences in gaze patterns [93]. Moreover, studies found that paradigms eliciting the N400 effect concurrently cause pupil size modulations [94,95], a response we already exploited in hybrid approaches for error detection [96,97]. Such considerations should be the subject of future works, as elevating classification accuracy is a primary objective for practical BCIs.

## 5. Conclusion

In this work, we designed an interactive paradigm that resembles a common use case for AR technologies, i.e., providing on-demand access to complementary situated information about physical entities. As visualizations of such information draw from various means of data representations, the presented task consequently featured both symbolic and non-symbolic stimuli. This study advances the characterizations of the neurophysiological activity associated with processing incongruent information in AR, revealing N400 and P600 components. We further observed a delayed response to non-symbolic numbers possibly related to the increased demands in processing information using more complex visualizations. Decoding of incongruency processing for BCIs yielded comparable performance for the implemented classification methods sLDA, SVM, and EEGNet. Future research should address open challenges, such as improving the classification accuracy of incongruency decoders, which is currently below the necessary for practical applications, and their implementation in online scenarios.

## CRediT authorship contribution statement

**Michael Wimmer:** Conceptualization, Methodology, Formal Analysis, Investigation, Writing - Original Draft, Writing - Review & Editing, Visualization. **Alex Pepicelli:** Methodology, Software. **Ben Volmer:** Investigation, Writing - Review & Editing. **Neven ElSayed:** Conceptualization, Writing - Review & Editing. **Andrew Cunningham:** Conceptualization, Resources, Writing - Review & Editing. **Bruce H. Thomas:** Conceptualization, Writing - Review & Editing. **Gernot. R. Müller-Putz:** Conceptualization, Writing - Review & Editing, Supervision. **Eduardo E. Veas:** Conceptualization, Writing - Review & Editing, Supervision, Funding Acquisition.

## Acknowledgments

The Know Center is funded within the Austrian COMET Program - Competence Centers for Excellent Technologies - under the auspices of the Austrian Federal Ministry of Transport, Innovation and Technology, the Austrian Federal Ministry of Economy, Family and Youth, and by the State of Styria. COMET is managed by the Austrian Research Promotion Agency FFG. This work was supported by the TU Graz Open Access Publishing Fund.

## Notes

### Competing Interest Statement

The authors have declared no competing interest.

